# Using confidence inferred from pupil-size to dissect perceptual task-strategy: support for a bounded decision-formation process

**DOI:** 10.1101/269159

**Authors:** Katsuhisa Kawaguchi, Stephane Clery, Paria Pourriahi, Lenka Seillier, Ralf Haefner, Hendrikje Nienborg

## Abstract

During perceptual decisions subjects often rely more strongly on early rather than late sensory evidence even in tasks when both are equally informative about the correct decision. This early psychophysical weighting has been explained by an integration-to-bound decision process, in which the stimulus is ignored after the accumulated evidence reaches a certain bound, or confidence level. Here, we derive predictions about how the average temporal weighting of the evidence depends on a subject’s decision-confidence in this model. To test these predictions empirically, we devised a method to infer decision-confidence from pupil size in monkeys performing a disparity discrimination task. Our animals’ data confirmed the integration-to-bound predictions, with different internal decision-bounds accounting for differences between animals. However, the data could not be explained by two alternative accounts for early psychophysical weighting: attractor dynamics either within the decision area or due to feedback to sensory areas, or a feedforward account due to neuronal response adaptation. This approach also opens the door to using confidence more broadly when studying the neural basis of decision-making.

## Introduction

During perceptual discrimination tasks subjects often rely more strongly on early rather than late sensory evidence even when both are equally informative about the correct decision e.g. ^1–4^. (But note that some studies in rodents and humans reported uniform weighting of the stimulus throughout the trial ^5–7^). From the perspective of maximizing the sensory information and hence performance such early weighting is non-optimal. Understanding this behavior may shed light on how the activity, or the read-out of sensory neurons limits our perceptual abilities, a major goal of contemporary neuroscience (e.g. ^8–10^). The classical explanation for such early psychophysical weighting is that it reflects an integration-to-bound decision-process in which sensory evidence is ignored once an internal decision-bound is reached ^1^. For simple perceptual discrimination tasks, decision confidence can be defined statistically ^11^, and hence also measured for such a model. Here, we derived new predictions of this model for how the temporal weighting of sensory evidence should vary as a function of decision confidence on individual trials. These revealed characteristic differences in the temporal weighting for high and low confidence trials, depending on the decision bound. We then sought to test these predictions in macaques performing a fixed duration visual discrimination task while also measuring the animal’s subjective decision confidence.

Measuring decision confidence psychophysically is relatively difficult, particularly in animals, and increases the complexity of a task, as e.g. for post-decision wagering ^12,13^, hence requiring additional training. To avoid these difficulties we devised a metric based on the monkeys’ pupil size. Combining this metric for decision confidence with psychophysical reverse correlation ^3,14,15^ allowed us to quantify the animals’ psychophysical weighting strategy for different levels of inferred decision-confidence, and test our model predictions. The animals showed clear early psychophysical weighting on average. But separating this analysis by inferred decision confidence revealed that early psychophysical weighting was largely restricted to high confidence trials. In fact, on low inferred confidence trials the animals weighted the stimulus relatively uniformly or even slightly more towards the end of the trial. Such behavior matched the predictions of the integration-to-bound model. Furthermore, the differences between both animals could be accounted for by the model by differences in the only free parameter – their internal decision-bound.

In contrast, the animals’ behavior could not be fully explained by two alternative accounts of early psychophysical weighting. The first alternative account are models in which the decision-stage provides self-reinforcing feedback to the sensory neurons ^16^, as suggested, e.g. for probabilistic inference ^17^, or by attractor dynamics within the decision-making area ^28^. The second, recent alternative proposal is that the early weighting simply reflects the feed-forward effect of the dynamics (gain control or adaptation) of the activity of the sensory neurons ^4^. Although each of these alternatives predicts the early weighting, we were unable to fully capture the animals’ data with the temporal weighting predictions of these models when separating trials by decision-confidence.

Together, our data suggest that the animals rely on a bounded decision-formation process. In this model, evidence at the end of the trial is only ignored once a certain level of decision-confidence is reached, thereby reducing the impact on performance. Moreover, this combination of techniques provides a novel tool for a more fine-grained dissection of an animal’s psychophysical behavior.

## Results

### Integration-to-bound models predict characteristic differences in temporal sensory weighting when separating trials by decision-confidence

Subjects during psychophysical discrimination task often give more weight to the early than late part of the stimulus presentation even in tasks when both are equally informative about the correct answer ^1,3,4^. We refer to this behavior as early psychophysical weighting, and the standard computational account is that it reflects an integration-to-bound decision process ^1^. In brief, this explanation suggests that subjects accumulate sensory evidence only up to a predefined bound not only in reaction time tasks but also in tasks when the stimulus duration is fixed by the experimenter, and when a complete accumulation of evidence over the course of the entire trial would be optimal. As a result, sensory evidence after the internal bound is reached is ignored and, together with a variable time at which this bound is reached, *on average*, early evidence is weighted more strongly than evidence presented late in the trial. If this explanation for the observed early weighting is correct, then across trials in which the decision-variable never reaches the bound, all evidence would be weighted equally, regardless when it was presented during the trial.

Interestingly, for simple perceptual discriminations tasks, decision confidence can be defined statistically ^11^, and directly linked to the decision-variable. In an integration-to-bound model it simply reflects the distance of the decision-variable to the category boundary. Here, we exploited this link and systematically explored how the temporal weighting of the sensory stimulus should depend on decision-confidence according to the integration-to-bound model. To do so we categorized trials into high or low confidence trials (median split) and measured the temporal weighting of the sensory evidence as the amplitude of the psychophysical kernel (PKA) over time (see Methods) for each category. We compared these for high confidence trials, low confidence trials and across all trials while systematically varying the decision bound of the model (Fig. **1**). As expected we found that the average PKA decreases more steeply if the decision bound is lower (see black lines in Fig. **1a** through **1e**), indicating that the decision-bound was reached earlier on average, and therefore the sensory evidence ignored from an earlier point in the trial. It is also intuitive that the PKA was typically larger for high compared to low confidence trials reflecting the stronger sensory evidence, and hence confidence, on those trials. Note that if the decision-bound is low, the decision-bound is reached on a large proportion of trials, and the assigned decision-confidence identical. These trials are therefore randomly assigned to the high and low confidence category, resulting in the similarity of the PKAs (Fig. **1a**). However, an interesting, non-trivial characteristic emerges for intermediate values of the decision bound (Fig. **1b-c**). Relatively strong evidence early during the trial led to high-confidence and early reaching of the decision boundary, resulting in the pronounced decrease of the PKA for high confidence trials. But for low confidence trials, the PKA not only showed no decrease but an increase over time (Fig. **1b-d**). As a result the PKAs for high and low confidence trials crossed and the PKA for low confidence trials exceeded that for high confidence trials at the end of the stimulus presentation. Over a range of values of the decision-bound the difference between the PKA for high and low confidence trials was therefore negative (Fig. **1f**). This characteristic behavior was even more pronounced when we defined decision-confidence not only based on evidence but also decision time, as previously suggested ^12,18^ (cf. Fig. **1g-l**). (Since our analysis depended only on the rank-order of the decision confidence these results hold generally, regardless of the relative weighting of time and evidence for decision confidence, see Methods.) Note that after sorting zero-signal trials by decision-variable, the PKA cannot easily be interpreted as a weight on the stimulus. For instance, the temporal weights on any one trial are always a non-zero constant starting at the beginning of the trial, and zero after some point. As a result, the averaged weights across all trials must be decreasing. The fact that the PKA may be increasing is the result of sorting the trials by confidence which separates the stimulus distributions between the high and the low signal trials. Equally, the more pronounced early difference in PKAs for low decision bounds (cf. Fig. **1a** and **1g**) reflects the fact that when decision-confidence is based on both time and evidence, trials with stronger early sensory evidence, and hence early decision-times, are assigned to the high confidence category. Nonetheless, these simulations reveal characteristic predictions about how a particular statistic – the psychophysical kernel as measured by taking the difference between the choice-triggered averages – should vary as a function of confidence for a bounded decision-formation process. We therefore next aimed to test these predictions in monkeys performing a visual discrimination task for which early psychophysical weighting was previously reported ^3^.

**Figure 1.**
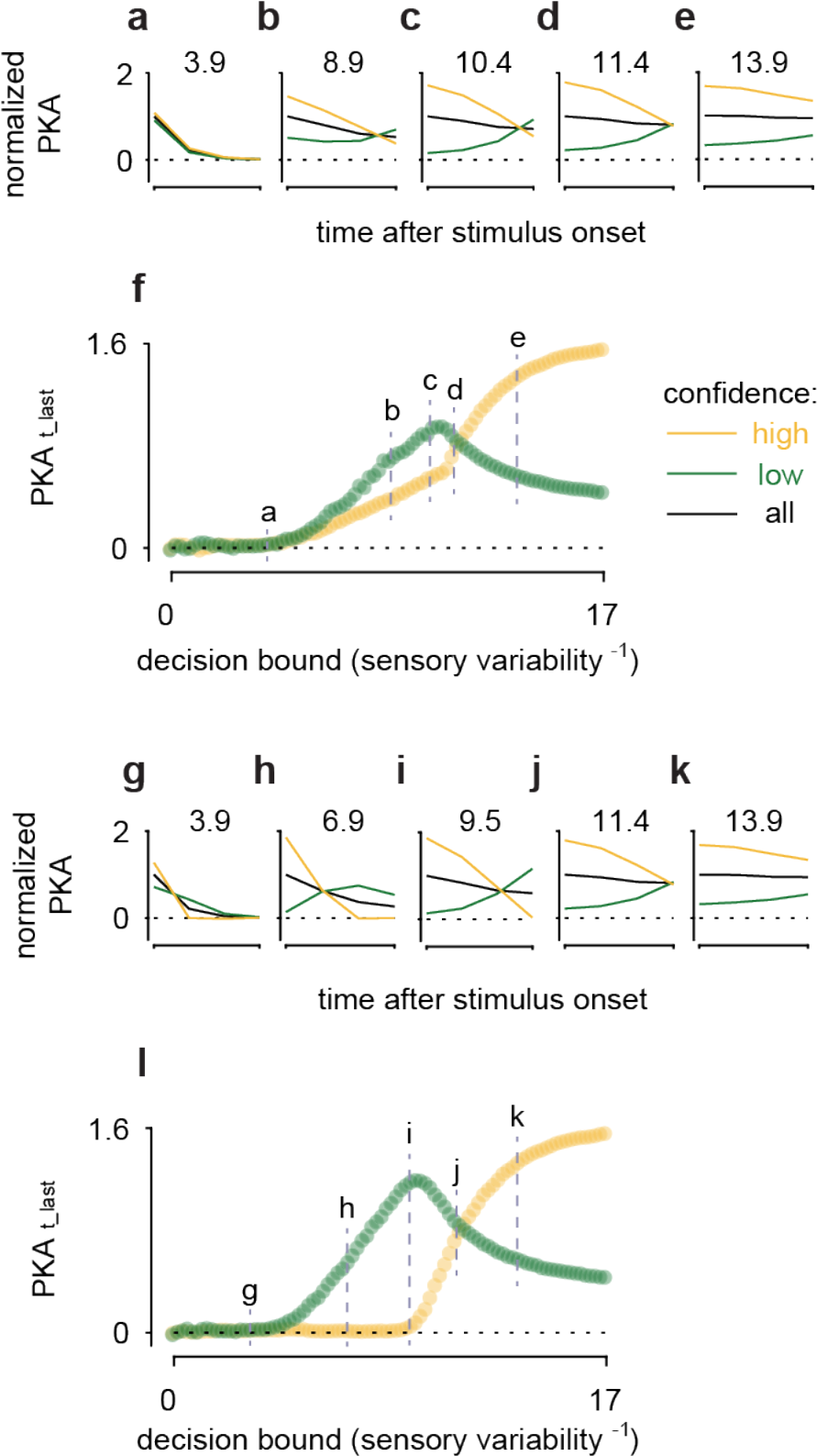
Integration-to-bound models predict characteristic differences in temporal sensory weighting for high and low confidence trials. **a-e)** The amplitude of the psychophysical kernel (PKA) is plotted over time for integration-to-bound models with different decision bounds. PKAs for low confidence, high confidence and averaged across all trials are shown in green, yellow and black, respectively, and normalized by the peak of the average psychophysical kernel. Note that for intermediate levels of the decision-bound the PKAs cross such that the PKA for low confidence trials exceeds that for high confidence trials at the end of the stimulus presentation. The value of the decision bound is marked in each panel. **f)** PKA_t_last_ is plotted for high (yellow) and low (green) confidence trials. The difference, ΔPKA_t_last_, depends characteristically on the level of the decision-bound in the model and the stimulus strength. Note that the decision-bound is normalized by the standard deviation of the sensory variability. The relationship between ΔPKA_t_last_ and the value of the decision bound therefore holds generally across tasks with different stimulus variability. **g-l)** Same as **a-f)** but for in an integration-to-bound model in which decision-confidence is based on both decision-time and evidence. Note that since our analysis only relied on the rank-order of the decision-confidence the results are independent of the relative weight of these influences on decision-confidence.

### The animals exhibit early psychophysical weighting behavior in this task

Two macaque monkeys performed a coarse disparity discrimination task (Fig. **2a**), similar to that described previously ^3^. The animals initiated each trial by fixating on a small fixation marker, and after a delay of 500ms a dynamic random dot stimulus was presented for a fixed duration of 1500ms. The stimulus was a circular random dot pattern defining a central disk and a surrounding annulus. The animals’ task was to determine whether the disparity-varying center was either protruding (“near”) or receding (“far”) relative to the surrounding annulus. Following the stimulus presentations two choice targets appeared above and below the fixation point, one symbolizing a “near” choice, the other a “far” choice. Importantly, the positions of the choice targets were randomized between trials such that the animals’ choices were independent of their saccade direction. While the animals performed this task we measured their eye positions and pupil size.

Similar to previous findings, e.g. ^1,3,4^ the animals relied more strongly on the stimulus early than late during the stimulus presentation. We quantified this as a decrease in the PKA (see Methods) throughout the stimulus presentation (Fig. **2b**). In order to test the model predictions separated by decision-confidence in the animals’ data we therefore sought to devise an approach to infer the animals’ decision confidence from pupil size measurements in this task.

**Figure 2.**
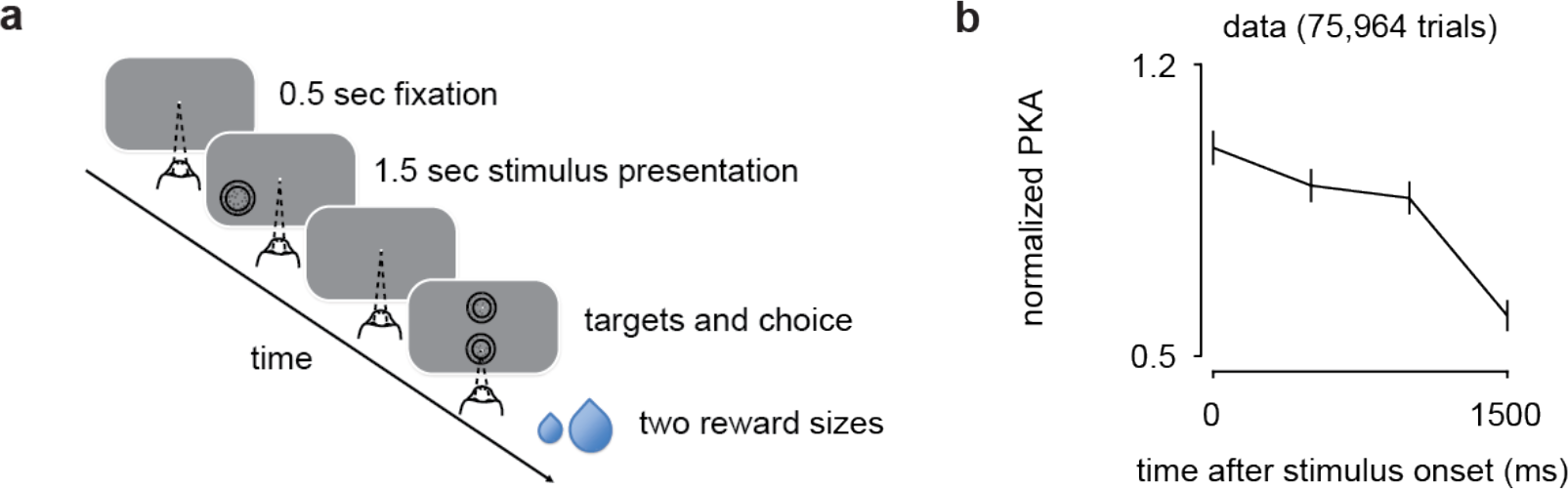
Task and early psychophysical weighting behavior. **a)** Two choice disparity discrimination task. After the animals maintained fixation for 0.5 sec the stimulus was shown for 1.5 sec. The animals had to decide whether the stimulus was ‘far’ or ‘near’ by making a saccade to one of two targets after the stimulus offset and received a liquid reward for correct choices, **b)** The time-course of the psychophysical kernel amplitude (normalized) shows that the animals weight the stimulus more strongly early during the trial. Data were obtained from 0% signal small available reward trials and collapsed across animals (A: 36,222 trials in 213 sessions, B: 13,334 trials in 84 sessions). Error bars are SEM derived by resampling.

### Pupil size is systematically associated with experimental covariates, consistent with pupil-linked changes in arousal

Pupil size has been linked to a subjects’ arousal in both humans ^19^ and monkeys ^20–23^. Our animals performed a substantial number of trials in each session (mean; animal A: 828, animal B: 1067). We therefore wondered whether a signature of their decreasing motivation with increased satiation during the behavioral session could be found in the animals’ pupil sizes. To this end we split the trials of each session into five equally sized bins (quintiles) and computed the average pupil size aligned on stimulus onset (Fig. **3a**). For these averages only 0% signal trials on which the available reward size was small (see Methods) were used. Moreover, to allow for the detection of slow trends throughout the session the pupil size data were not high-pass filtered for this analysis. We found that in both animals pupil size systematically decreased throughout the session, as expected for a decrease in arousal with decreased motivation or task engagement with progressive satiation.

We next explored the effect of varying the available reward size in a predictable way (see Methods). Consistent with previous results ^24^ the animals’ psychophysical performance on large available reward trials exceeded that on small available reward trials (Fig. **3d**). When averaging the time-course of the pupil size for 0% signal trials separated by available reward size, we found that pupil size for large available reward trials increased progressively compared to that on small available reward trials (Fig. **3b**). The animals were rewarded after correct choices following the stimulus presentation. The time-course of this pupil-size modulation with available reward size is therefore consistent with modulation related to the animals’ expectation of the reward towards the end of the trial. Indeed, the difference in mean pupil with available reward size over the last 250ms of the stimulus presentation was highly statistically reliable (Fig. **3e**), similar to previous findings ^25^.

Note that previous studies that revealed arousal linked pupil size modulation typically used long inter-trial-intervals lasting several seconds ^20–23^, which were deemed necessary to stabilize pupil size prior to stimulus or trial onset. Conversely, our task allowed for short inter-trial-intervals (animal A: 65-4772ms, median: 136ms; animal B: 115-3933,median: 146ms) to yield a large number of trials per session. Nonetheless, the above analyses revealed robust signatures of pupil size modulation with experimental manipulations of arousal also for this task.

Previous work in humans found that pupil size increased with task difficulty, which is thought to reflect changes in arousal related to “cognitive load” or “mental effort” ^26–28^. To explore whether such a signature was evident for our task, we divided our data into easy (≥50% signal) and hard trials (≤10% signal, excluding 0% signal trials) (Fig. **3c**). To remove effects of available reward size this analysis was restricted to small available reward trials. Consistent with the expected modulation for cognitive load, pupil size in hard trials weakly exceeded that for easy trials in the initial period of the stimulus presentation (before ~750ms after stimulus onset). However, the more pronounced modulation with task difficulty occurred in the opposite direction towards the end of the trial.

**Figure 3.**
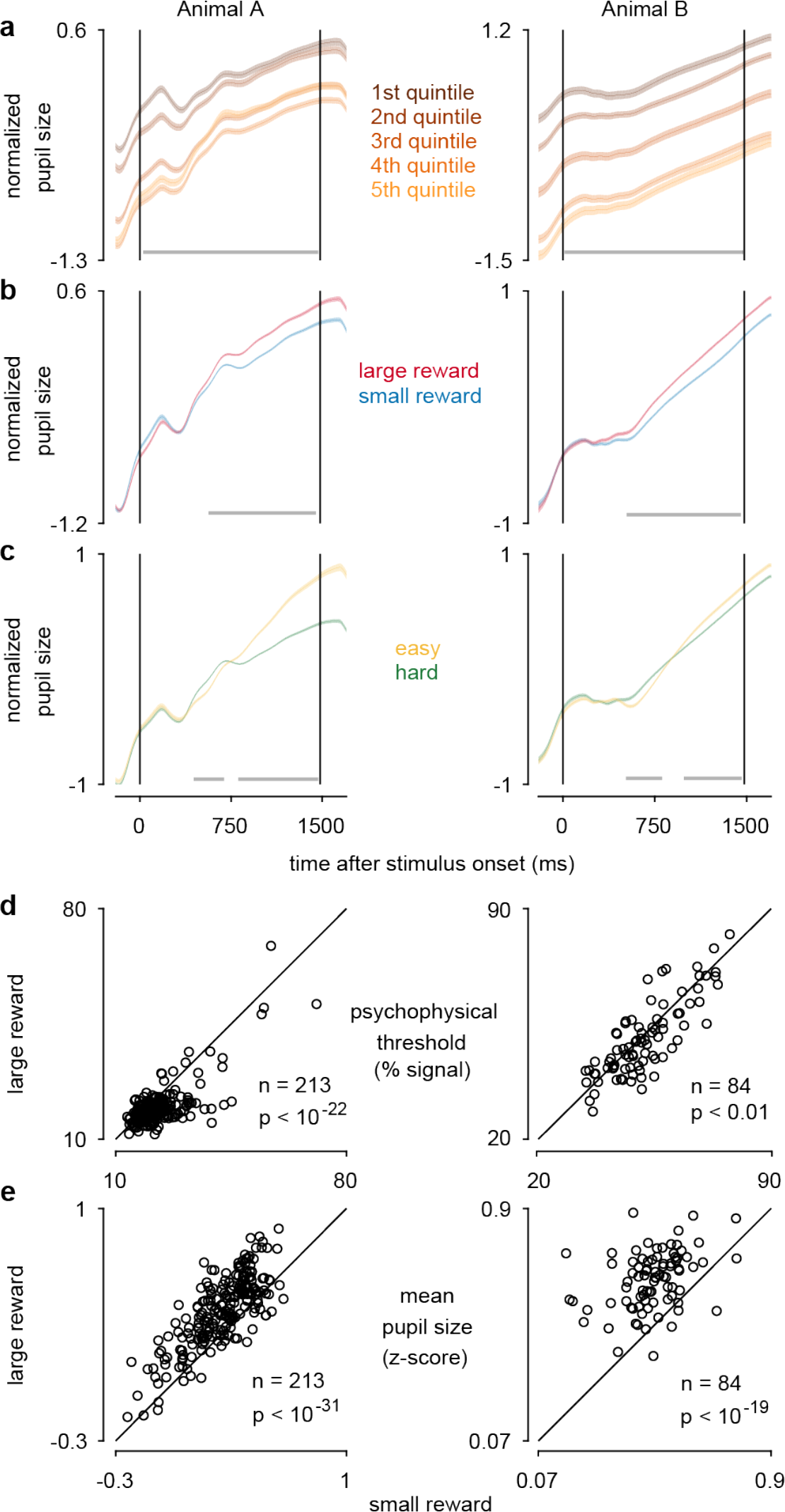
Pupil size modulation with task covariates is consistent with pupil-linked arousal. **a-c** Average z-scores (across conditions) ± SEM of pupil size aligned on stimulus onset are shown for monkey A (left) and B (right). Horizontal lines at the bottom of each panel depict epochs of significant (p<0.05, corrected for multiple comparisons) pupil size modulation (by ANOVA in **a)**, two sample t-tests in **b-c)**. **a)** Mean pupil size for five equally sized bins throughout each experimental session. Only small available reward 0% signal trials are used. Pupil size decreases throughout the session as expected for decreasing motivation. (A: 6,987 trials from 213 sessions, B: 2,571 trials from 84 sessions). **b)** Average time courses of pupil size on 0% signal trials for large (red) and small (blue) available reward trials. (A: 18,855 small available reward trials and 18,678 large available reward trials from 213 sessions, B: 6,843 small available reward trials and 6,832 large available reward trials from 84 sessions) **c)** Average time courses of pupil size on hard (<10%, excluding 0% signal, green) and easy (≥50% signal, yellow) trials. Only trials with the small available reward were used. (A: 39,390 hard trials and 8,651 easy trials from 213 sessions, B: 10,813 hard trials and 14,020 easy trials from 84 sessions). **d)** Psychophysical thresholds on high available reward trials were significantly smaller than in small available reward trials (A: n = 213, p < 10^−22^, B: n = 84, p < 0.01). **e)** Average pupil size during the 250ms prior to the stimulus offset were significantly larger in large compared to small available reward trials in trials (A: n = 213, p < 10^−31^, B: n = 84, p < 10^−19^, all paired t-tests).

Remarkably, plotting this modulation across training sessions revealed that this late modulation only emerged once the animals knew the task well (Fig. **4a**) and was correlated with task performance (Fig. **4b**). This late modulation appears to reflect the animals’ expectation to receive a reward based on their knowledge of the probability of being correct given the stimulus difficulty. It might thus be interpretable as a modulation based on the animal’s confidence to make the correct decision. We will show next that this modulation indeed exhibits established key signatures ^11,29^ of decision confidence, supporting this interpretation.

**Figure 4.**
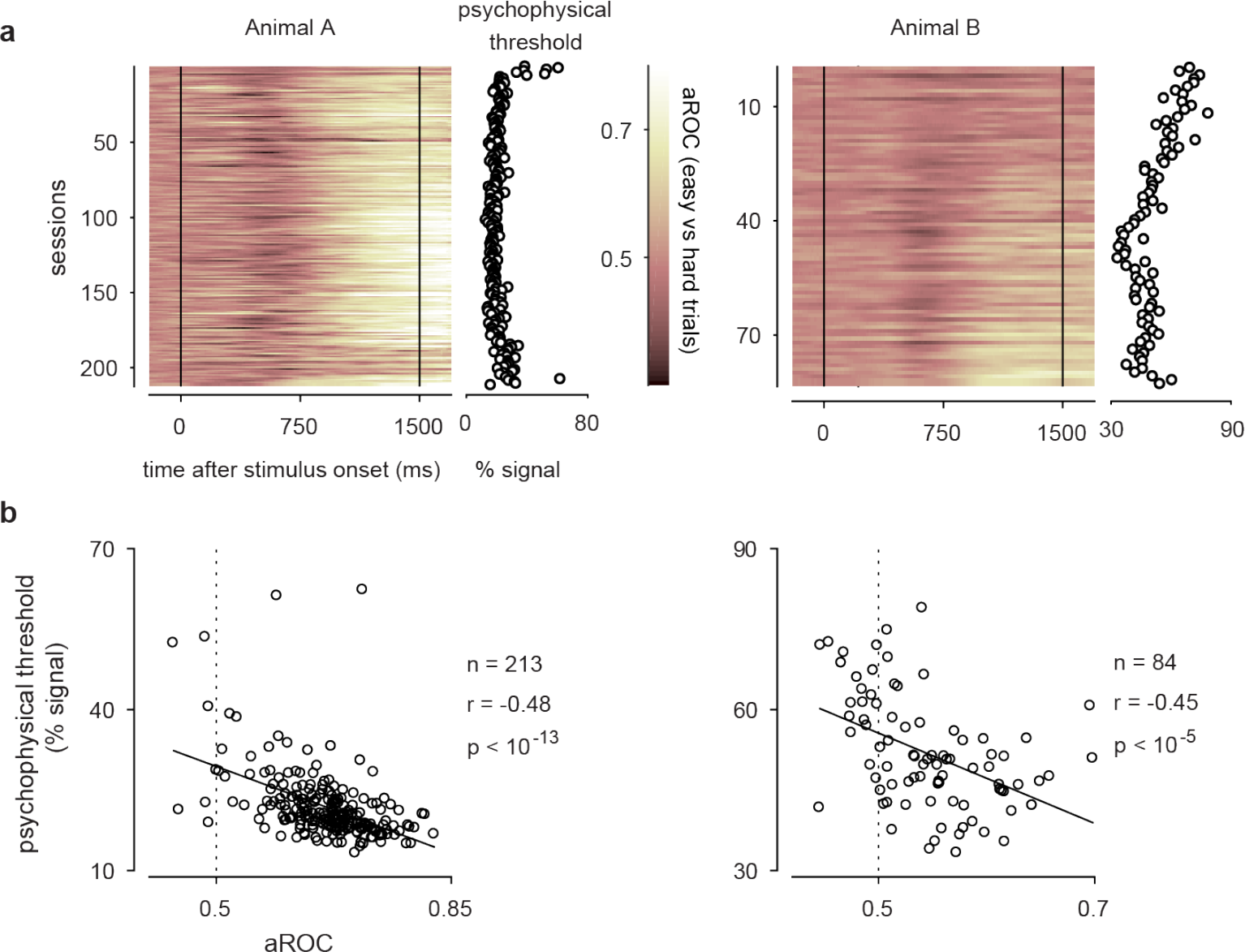
The signature of decision-confidence requires good task performance. **a)** Discriminability between hard (<10%, excluding the 0% signal) and easy (≥50% signal) trials, quantified as aROC for each session (ordinate; 213 sessions from animal A, 84 sessions from animal B), plotted as a function of time (abscissa) in the trial after stimulus onset. Note that the systematically larger pupil size for easy trials (bright colors) late in the trial emerge only after extensive training, particularly in monkey B. **b)** The average aROC during the 250ms prior to the stimulus offset is significantly correlated with the psychophysical threshold (A: n = 213, r = −0.48, p < 10^−13^, B: n = 84, r = −0.45, p < 10^−5^; Pearson’s correlation coefficient).

### Pupil size in this task can be used to infer the animal’s decision confidence

For a two-alternative sensory discrimination task analogous to the one used here decision confidence is monotonically related to the distance to a category boundary ^11,30^, i.e. the integrated sensory evidence, as schematically shown in Fig. **5a**. From a statistical perspective decision confidence in such discrimination tasks should be systematically associated with evidence discriminability, accuracy and choice outcome (model predictions in Fig. **5b** top row). Empirically, we found that mean pupil size during the 250ms before stimulus offset showed the three characteristics of statistical decision confidence keeping reward size constant (we restricted these analyses to small available reward trials to eliminate the effect of available reward size). The findings were qualitatively the same when only analyzing large available reward trials (supplementary Fig. **1**). First, in both animals, pupil size increased monotonically with performance accuracy (Fig. **5b**, first column). Second, when separating trials based on pupil size (median split), the animals showed better discrimination performance for trials on which pupil size was larger, as expected for improved evidence discrimination with higher decision confidence ^11^ (Fig. **5b**, middle column). Third, as predicted, when separating correct and error trials, decision confidence increased on correct and decreased on error trials. Interestingly, we also observe a slight increase in pupil size with signal strength for higher signal strengths in animal B. Such a pattern is expected if decision confidence is informed not only by the strength of the sensory evidence, as described above, but also by decision time as observed in human observers ^18^.

Since we used a white fixation marker our results pupil size measurements might in principle have been affected by the animals’ fixation precision. To control for this potential confound we therefore performed a number of control sessions in which instead of a white fixation dot we used a black fixation marker. If our results were mostly driven by differences in luminance resulting from differences in fixation precision across conditions the modulation with our experimental co-variates should reverse. However, our results were robust when instead of a white fixation marker we used a black fixation marker (see supplementary Fig. **2**). Together, these analyses support our conclusion that mean pupil size at the end of the stimulus presentation can be used to infer the animals’ decision confidence.

**Figure 5.**
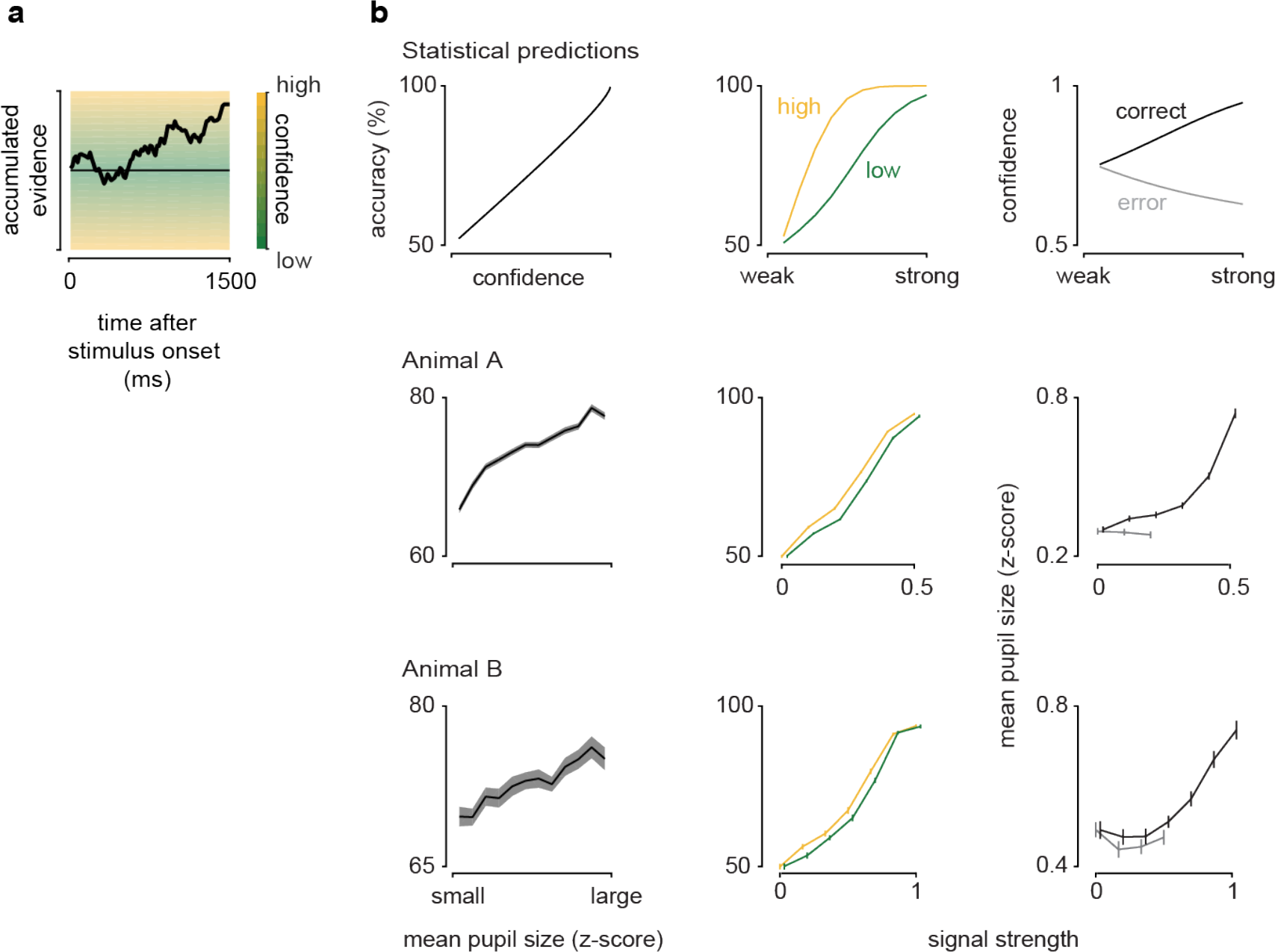
Pupil size shows signatures of decision confidence. **a)** Schematic of a drift-diffusion model in which the decision confidence depends on the distance of the decision variable to the category boundary. **b)** Signatures of statistical decision confidence (top row) are compared to our metric based on pupil size (average pupil size during the 250ms prior to stimulus offset) (middle and bottom rows). Left column: Statistical decision confidence predicts accuracy. Similarly, mean pupil size increases monotonically with accuracy. Middle column: For high decision confidence statistical decision confidence predicts steeper psychometric functions than for low decision confidence. The monkeys’ psychometric functions separated by mean pupil size are slightly steeper for large compared to small mean pupil size, as predicted for decision confidence. Right column: Decision confidence is predicted to increase with signal strength in correct trials and decrease with signal strength in error trials. Mean pupil size increases for correct and slightly decreases on error trials (monkey A), and for low signal strengths in monkey B. Data points are slightly offset for better visualization. For animal A all the sessions were included (213 sessions). For animal B analyses are restricted to the last 40 sessions with good performance (cf. Fig. **4**). Data are shown as mean ± SEM.

### The animals’ data separated by inferred decision confidence supports the predictions of the integration-to-bound model

Having established the relationship between pupil-size and decision confidence in our task, we now use it to test the confidence-related predictions of the integration-to-bound model using our data. To do so, we computed the animals’ psychophysical kernels separately after categorizing high or low inferred decision confidence trials (median split based on the pupil-size metric). For inferred high-confidence trials, we observed a decrease in psychophysical kernel amplitude (PKA) for both monkeys. In contrast, for inferred low confidence trials the PKA either stayed relatively constant throughout the trial (monkey B, Fig. **6c**), or first increased and then decreased (monkey A, Fig. **6b**). Furthermore, the PKA at the end of low-confidence trials was approximately equal (monkey B) or higher (monkey A) than the PKA for high-confidence trials. Importantly, the data for both monkeys best agree with the predictions of an integration-to-bound model when subjective confidence is based on both evidence and time ^18^ with the difference between the two animals naturally explainable by differing internal integration bounds (cf Fig. **1i** and **1j**).

We next wondered whether the data was also explainable by two alternative accounts of the early psychophysical weighting: first, models with attractor dynamics resulting from recurrent feedback, or second a purely feed-forward account that includes adaptation.

**Figure 6.**
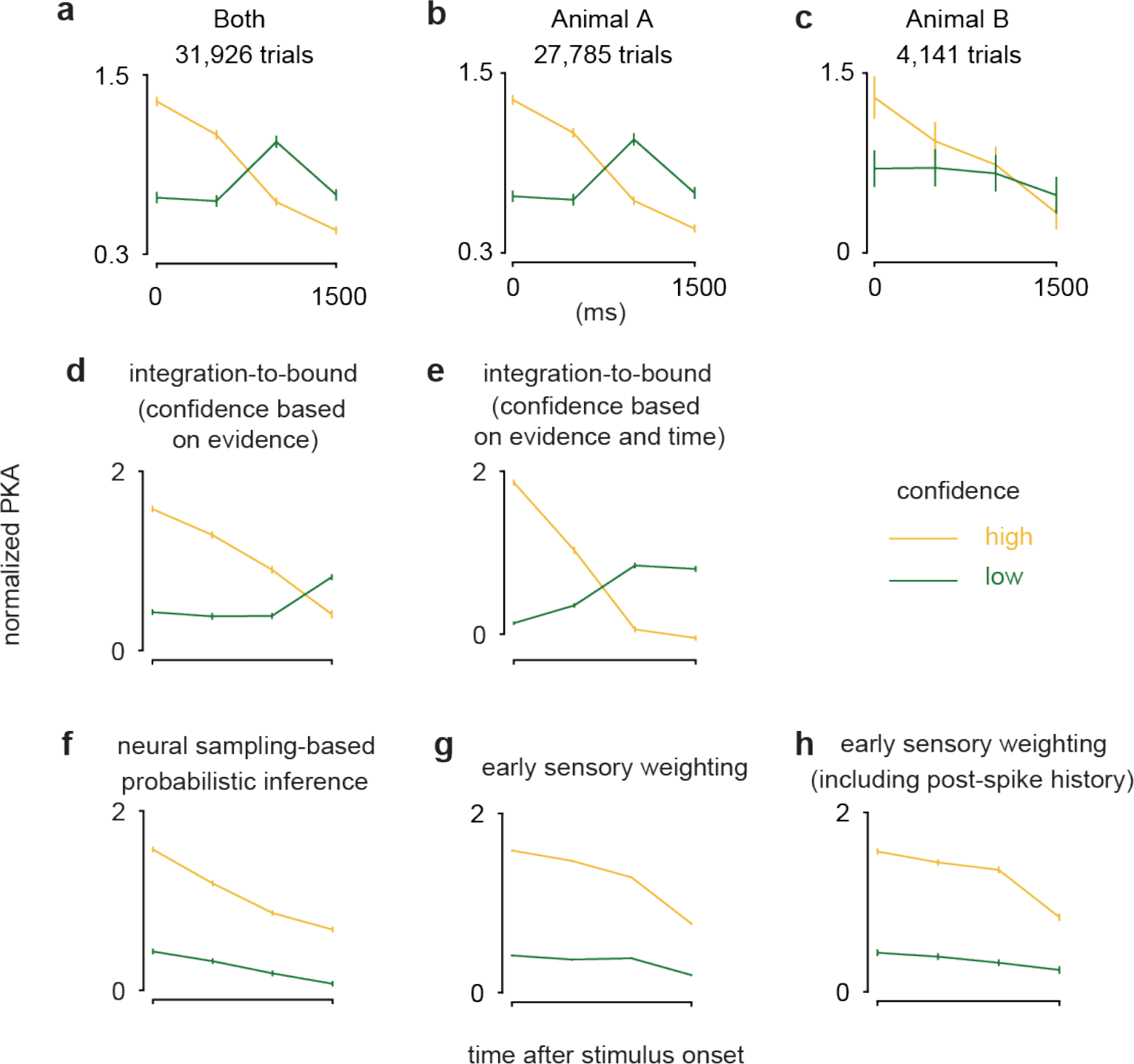
The animals’ psychophysical weighting on low and high confidence trials is compared to model predictions. Psychophysical kernel amplitudes for high (yellow) and low (green) confidence trials (median split) are plotted as a function of time. **a)** Psychophysical kernel separated by confidence inferred from pupil size. Data from 0% signal trials were collapsed across animals **a)** and shown separately for each animal **(b, c)** (A: 213 sessions, B: 40 sessions. To avoid confounding the pupil size modulation for available reward size with that for inferred decision-confidence, the median split based on pupil size to assign trials to the high or low confidence bin was performed separately for small and high available reward trials.) Note the similarity of this result to the prediction by an integration-to-bound model (Fig. **6d**, **e**). **d)** Integration-to-bound model in which trials were separated based on decision confidence defined as |decision variable|. **e)** Integration-to-bound model in with decision confidence depended on both |decision variable| and the model’s decision time on each trials (see Methods). **f)** Neural sampling-based probabilistic inference model for which decision-confidence is defined by the Bayesian posterior probability. **g)** Early sensory weighting model after ^4^ based on a linear-nonlinear model reflecting the response dynamics (gain control) of sensory neurons. **h)** An extension of the model used in **g)** to also include a post-spike filter to capture a neuron’s spiking history^4^. Error bars (SEM) were derived by resampling.

To test the first alternative account, we implemented a model ^17^ in which the decrease of the amplitude of the psychophysical kernel results from self-reinforcing feedback from decision neurons to sensory neurons. Because of its recurrent connectivity this model exhibits attractor dynamics, in which early evidence is effectively weighted more strongly than evidence presented late in the trial. Other recurrent models of perceptual-decision making, whether across cortical hierarchies ^16^, or proposing attractor dynamics within the decision area itself ^31,32^ share this attractor behavior. In these models the behavior of decision variable after stimulus onset can be described by a double-well energy landscape, where the minimum of each well corresponds to a choice attractor (cf. ^16^; inset in their Fig. **2d**). As a result, the effect of early evidence on the decision variable will be amplified by the subsequent pull exerted by whatever attractor towards which the early evidence had pushed the decision variable. While this behavior resembles that of the integration-to-bound model, it differs in its predictions when separating trials according to confidence (Fig. **6f**). Specifically, we were unable to identify model parameters for which the kernel amplitude in low confidence trials exceeded that for high confidence trials at the end of the stimulus presentation (supplementary Fig. **4a**). In order to convince ourselves that an attractor dynamic by itself is indeed unable to account for our data, we confirmed this finding for two idealized attractor models in which attractor strength and hence slope of the PKA were determined by a single parameter (similar to the integration-to-bound model) - see Supplementary Fig. **4b-c**. As for the neural sampling-based probabilistic inference model, varying this parameter did not yield kernels for which the kernel amplitude in low confidence trials exceeded that for high confidence trials at the end of the stimulus presentation. Indeed, the only way to achieve a similar late-trial PKA for high and low confidence was to strengthen the attractor dynamics in one of the models to a degree that made the late-trial PKA close to zero - in contradiction to the data (see supplementary Fig. **4b** for details).

Finally, we tested the behavior of two versions of an early sensory weighting model after ^4^ (their Fig. **4a** and **6a**), in which the decrease in PKA results from adaptation of the sensory responses in a purely feed-forward way. The model generates choices based on the integrated inputs of stimulus-selective sensory neurons, whose response decreases over the time of the stimulus presentation. Such decrease in response amplitude after response onset is typically observed for sensory neurons and may reflect a gain control mechanism or stimulus-dependent adaptation. As expected, we found a decreasing PKA across all trials. But like for the attractor-based models investigated above, and unlike for our data, the amplitude of the high-confidence PKA was consistently larger than the low-confidence PKA (Fig. **6g**). This pattern remained unchanged over a wide range of model parameters that yielded plausible sensory responses (compare supplementary Fig. **4d**). We also extended this model to include a post-spike filter ^4^ to account for a neuron’s refractory period and autocorrelation of the spiking response (Fig. **6h**). Similar to the model without the post-spike filter, the amplitude of the psychophysical kernel for high confidence trials was consistently higher than that for low confidence trials, differing from the animals’ behavioral data.

Together, these results indicate that while each of these models could account for early psychophysical weighting, a decision bound was necessary to account for the monkeys’ behavioral differences with inferred decision-confidence.

## Discussion

The frequently observed ^1–4^ early weighting of sensory evidence in perceptual decision-making tasks has classically been explained to reflect an integration-to-bound decision process ^1,33^. Here, we first derived decision confidence-specific predictions for this account. Second, in order to test these predictions, we devised a metric based on pupil size that allowed us to estimate two macaques’ subjective decision confidence on individual trials without the use of a wagering paradigm. Finally, we compared our confidence-specific data to two alternative accounts of early weighting – attractor dynamics and response adaptation – and found that neither of those models could explain our data. This combined approach provided new insights into the animals’ decision-formation process. It revealed that the frequently observed ^1–4^ early weighting of the sensory evidence was largely restricted to high-confidence trials, and that the shape of the psychophysical kernel amplitude (PKA) confirmed our predictions based on the integration-to-bound model. In fact, the match between data and model was best when we incorporated a recent proposal about how subjective confidence was not just based on the strength of the presented evidence, but also integration time ^18^. Moreover, our data could not be fully explained by other computational accounts for early psychophysical weighting such as sensory adaptation ^4^ or models of perceptual decision-making with recurrent processing ^l6,17,32^. We note that our findings do not preclude the contribution of these alternative models. However, our results highlight that none of these accounts is sufficient to explain the data by itself and that a decision-rule that implements an early stopping of the evidence integration process appears necessary.

Our analysis of pupil size showed that even without the stabilizing effect of long inter-trial intervals pupil size was reliably correlated with experimental covariates, and could be used to infer the animal’s decision confidence. The correlation of pupil size with decision confidence is similar to that in a recent psychophysical study in humans ^34^ that queried decision confidence directly. As we did here, this study found a positive correlation between subjects’ pupil size before they made their judgment and their reported decision confidence. Previous work inferring an animal’s decision confidence typically relied on behavioral measurements such as postdecision wagering ^12,13^ and the time an animal is willing to wait for a reward ^35^, which increases the complexity of the behavioral paradigm and hence the required training of the animals. To our knowledge the present study is the first to relate pupil size measurements in animals to decision-confidence. Such a pupil-size based metric opens up studies of decision making in animals to include decision confidence without increasing the complexity of the behavioral paradigm.

In our task the animals were rewarded on each trial directly after making their choice. Consistent with modulation of pupil-linked arousal due to reward expectation ^25,36^, pupil size was progressively larger towards the end of the trial when the (known) available reward was large compared to when it was small (cf. Fig. **3b**). Such reward-based interpretation of the pupil-size modulation associated with decision-confidence may explain our and ^34^ findings here, which contrasts with studies associating increases in pupil size with uncertainty e.g. ^29,37–40^. Specifically, a recent study ^29^ observed the opposite relationship between inferred decision confidence and pupil size, measured after the subject’s perceptual report: larger pupil size after the subject’s report, and before receiving feedback, was associated with higher decision uncertainty. Access to information, e.g. whether or not a choice is correct, can be rewarding by itself ^41,42^. It may therefore be that in ^29^ the reward was such access to information, i.e. the feedback on each trial. When the confidence about the correct choice is low, the information is more valuable, hence resulting in the observed negative correlation with pupil size. Alternatively, this discrepancy may also reflect methodological differences such as the time-interval used for the analysis (before or after the choice was made, but see also ^38^). More generally, these findings underscore the importance to consider a subject’s motivational context when interpreting pupil size modulation.

Moreover, pupil-size modulation by cognitive factors has been linked to a number of neural circuits mirroring the complexity of the signal. These include the locus coeruleus noradrenergic system ^43,44^, a brain-wide neuromodulatory system involved in arousal, the inferior and superior colliculi, which mediate a subject’s orienting response to salient stimuli ^45,46^, but there is also evidence for an association with cholinergic modulation ^47,48^, which is also linked to attention.

The emergence of a reliable signature of decision-confidence required that the animals performed the task well (cf. Fig. **4**). We propose two possible, not mutually exclusive, accounts for this. First, in line with the notion that the observed pupil-size modulation linked to decision confidence is driven in part by reward expectation, it may reflect the animal’s improved knowledge of the timing of the task and hence the anticipation of the reward. Second, it may reflect the fact that in order to engage the pupil-linked arousal circuitry a certain threshold of decision-confidence needs to be exceeded. Such an interpretation would mean that once the signature of decision-confidence emerges a higher level of decision-confidence is reached at least on some trials.

Our animals’ psychophysical behavior separated by inferred decision-confidence was well described by a bounded accumulation decision process. These results imply that in a subset of trials sensory evidence was ignored after a certain level of decision-confidence had been gained. We find that in our task, across all difficulty levels, the loss in performance is small for the bounds required to explain our data (suppl. Fig. **5**). Since the overall loss will differ between different experiments, it might explain some of the differences seen in the temporal profile of PKAs across studies (e.g. ^1–5,7,49^). Furthermore, under the assumption that evidence accumulation is costly, it may provide a normative reason for the early termination of evidence integration ^50,51^.

## Materials & Methods

### Animal preparation and surgery

All experimental protocols were approved by the local authorities (Regierungspräsidium Tübingen). Two adult male rhesus monkeys (Macaca mulatta); A (7 kg; 11 years old) and animal B (8 kg; 11 years old), housed in pairs, participated in the experiments. The animals were surgically implanted with a titanium head post under general anesthesia using aseptic techniques as described previously ^52^.

### Visual discrimination task

The animals were trained to perform a two choice disparity discrimination task (Fig.**2a**). The animals initiated trials with the visual fixation on a small white fixation spot (size: 0.08-0.12°) located on the center of the screen. After the animals maintained fixation for 500ms, a visual stimulus was presented (median eccentricity for Animal A: 5.3°; range 3.0 – 9.0°, median eccentricity for Animal B: 3.0°, range 2.3 – 5.0°) for 1,500ms. After that two choice targets, each consisting of a symbol representing either a near or a far choice and whose positions were randomized between trials, appeared above and below the fixation spot. Once the fixation spot disappeared, the animals were allowed to make a choice via saccade to one of these targets. The animals received a liquid reward for correct choices. Randomizing target positions allowed us to disentangle saccade direction and choice.

### Visual stimuli

Visual stimuli (luminance linearized) were back-projected on a screen using a DLP LED Propixx projector (ViewPixx; run at 100Hz; 1920 × 1080 pixel resolution, 30 cd/m^2^ mean luminance) and an active circular polarizer (Depth Q; 200Hz) for animal B (viewing distance 97.5cm), or two projection design projectors (F21 DLP; 60Hz; 1920 × 1080 pixel resolution, 225 cd/m^2^ mean luminance, and a viewing distance of 149 cm) and passive linear polarizing filters for animal A. The animals viewed the screen through passive circular (animal A) or linear (animal B), respectively, polarizing filter. Stimuli were generated with custom written software using Matlab (Mathworks, USA) and the psychophysics toolbox ^53–55^.

The stimuli were circular dynamic random dot stereograms (RDS), which consisted of equal numbers of white and black dots, similar to those previously used ^3^. Each RDS had a disparity-varying circular center (3° diameter) surrounded by an annulus (1° wide) shown at 0° disparity. On each video-frame, all center dots had the same disparity whose value was changed randomly on each video-frame according to the probability mass distribution set for the stimulus. For the 0% signal stimulus the disparity drawn from a uniform distribution (typically 11 values in 0.05° increments from −0.25° to 0.25°). The monkeys were rewarded randomly on half of the trials on 0% signal trials. These 0% signal trials were randomly interleaved with near disparity or far disparity signal trials. For each session, one near and one far disparity value was used to introduce disparity signal by increasing the probability of this disparity on each video frame during the stimulus presentation on this trial. The range of signal strengths was adjusted between sessions to manipulate task difficulty and encourage performance at psychophysical threshold. Typical added signal values were 3%, 6%, 12%, 25% and 50%.

The choice target symbols were random dot stereograms very similar to 100% signal stimuli except that their diameter was smaller (2.2°).

To allow for constant mean luminance across the screen, equal numbers of black and white dots were used for the stimulus and the target symbols. Since we used a white fixation dot systematic differences in fixation precision could- in principle- influence our findings. If this were the case a black fixation marker should give the opposite results. We therefore also conducted control experiments using a black fixation marker, which yielded very similar results, indicating that systematic differences in fixation precision are insufficient to explain our findings.

### Reward size

To discourage the animals from guessing the available reward size was increased based on their task performance. After 3 consecutive trials with correct choices, the available reward size was doubled compared to the original reward size. After 4 consecutive trials with correct choices, the available reward size was again doubled (quadruple compared to the original size) and remained at this size until the next error. After every error trial, the available reward size was reset to the original.

### Pupil data acquisition and analysis

During the experiments, the animals’ eye positions and pupil size were measured at 500Hz using an infrared video-based eye tracker (Eyelink 1000, SR Research Ltd, Canada), digitized and stored for the subsequent offline analysis. The eye tracker was mounted in a fixed position on the primate chair to minimize variability in pupil size measurements between sessions. Our pupil analysis focused on the period of animals’ fixation in which the gaze angles were constant.

Only successfully completed trials (correct and error trials) were included for the analysis. During pre-processing we first down-sampled the pupil size data such that the sampling rate matched the refresh rates of our screens (60Hz for animal A, 100Hz for animal B), effectively low-pass filtering the data. We next high-pass filtered the data by subtracting on each trial the mean pupil size of the preceding 10 and following 10 trials (excluding the value of the current trial). This analysis removed linear trends on the pupil size within a session and was omitted for the analysis of pupil size changes throughout a session (Fig. **3a**). Finally, pupil size measurements were z-scored using the mean and standard deviation during the stimulus presentation period across all trials.

When comparing pupil size across conditions we aimed to minimize any mean difference of pupil size between conditions at stimulus onset. To do so, we computed a baseline pupil size, which was defined as the average pupil size in the epoch 200ms prior to stimulus onset, and iteratively excluded trials in which the baseline value deviated most from the condition with the higher number of trials until the absolute mean difference of the z-score of the baseline pupil size was below 0.05. This procedure successfully made the baseline pupil size statistically indistinguishable across conditions with a small loss of trials in each session (mean ± SD of the lost trials; Animal A, 6.89± 3.90%; Animal B, 8.24 ± 3.20%).

### Psychometric threshold

The animals’ choice-behaviors were summarized as a psychometric function by plotting the percentage of ‘far’ choices as a function of the signed disparity signals and then fitted with a cumulative Gaussian function using maximum likelihood estimation. The standard deviation of the cumulative Gaussian fit was defined as the psychophysical threshold and corresponds to the 84% correct level. The mean of the cumulative Gaussian quantified the subject’s bias.

### Psychophysical kernel

Psychophysical kernels were computed to quantify how the animals used the stimulus for their choices ^3,15^. Only 0% signal trials were used for this analysis. First, the stimulus was converted into an n-by-m matrix (n: number of discrete disparity values used for the stimulus; m: number of trials) that contained the number of video frames on which each disparity was presented per trial. Next, the trials were divided into ‘far’ choice and ‘near’ choice trials. The time-averaged psychophysical kernel was then computed as the difference between the mean matrix for ‘near’ choice trials and that for ‘far’ choice trials. We also computed a time-resolved psychophysical kernel as the psychophysical kernels for four non-overlapping consecutive time bins (each of 375ms duration) during the stimulus presentation period. Kernels were averaged across sessions, weighted by the number of trials in that session. The amplitude of the psychophysical kernels over time was calculated as the inner product between the time-averaged psychophysical kernel and the psychophysical kernel for each time bin. Kernel amplitudes separated by inferred decision confidence were then normalized by the maximum of the psychophysical kernel averaged across both conditions such that the relative differences between conditions remained. The standard error of the amplitude was calculated by bootstrapping (1000 repeats).

### Operationalizing decision-confidence

When viewed from a statistical perspective decision confidence can be linked to several behavioral metrics such as accuracy, discriminability and choices on error or correct trials ^11^ (Fig. **5b**). Here, we simulated an observer’s decision-variables on each trial analogously to ^29^. The decision variable (*d*) was drawn from a normal distribution whose mean depended on the signed signal strength (with negative and positive signal reflecting near and far stimuli, respectively) and the standard deviation on the observer’s internal noise (22.8 % signal, the median of the animals’ psychophysical thresholds). The sign of the *d* determined the choice on each trial. Assuming a category boundary c, trial-by-trial confidence (the distance between the decision variable and the category boundary) was transformed into a percent correct ^35^:

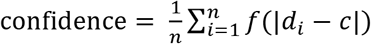

where f is the cumulative density function of the normal distribution.

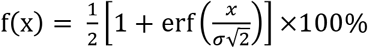

To simulate the relationship between accuracy and confidence, we generated the *d* for 10^8^ trials, binned these based on the level of confidence (20 bins) and computed the accuracy for each bin. To examine the relationship between confidence and psychophysical performance performed a median split of the trials based on confidence and measured the psychometric function for high and low confidence trials. Finally, we calculated the mean confidence as a function of signal strength separately for correct and error trials.

### Perceptual decision models

To compare the animals’ psychophysical kernels to different decision-strategies we simulated different perceptual decision models and calculated psychophysical kernels for the model data. For all simulations only 0% signal trials were used, and the model “decision-confidence” was defined as |decision-variable| at the end of each trial, unless stated otherwise. Psychophysical kernel amplitudes were then computed separately for high and low confidence trials, after a median split based on this metric for decision-confidence.

### Integration-to-bound model

In this model the decision-variable (*d*) is computed as the integrated time-varying difference of the population response of two pools of sensory neurons. (For the disparity discrimination task these would consist of one pool preferring near disparities, the other preferring far disparities.) We computed the time-varying population response as the dot product between the time-varying stimulus (analogous to that used in the experiments) and an idealized version of the animals’ time-averaged psychophysical kernel. On each trial, once the decision variable reached a decision bound (at decision time, *t*) ^1,33^ the decision-variable was fixed at that value (absorbing bound) until the end of the trial. The choice of the model was based on sign(*d*) at the end of the trial. We used two approaches to derive decision confidence for this model. First, it was defined as |d| at the end of the trial. This approach ignores the decision time. This model had one free parameter (the height of the decision bound), which we varied to best account for the time-courses of the psychophysical kernel amplitudes for low and high confidence trials. In this model, all trials in which the decision bound was reached are assigned the same confidence. Second, we also generated predictions for the proposal that subjective confidence is higher for those trials in which the bound is reached earlier ^12,18^. Since our analysis only relied on the rank-order of the trials based on confidence our results are independent of how exactly this time is converted into confidence.

### Neural sampling-based probabilistic inference model (Haefner et al 2016)

We used the model by ^17^, implemented for an orientation discrimination task. In this model, the decision is based on a belief over the correct decision (posterior-probability over the correct decision), which is updated throughout each trial. The decision-confidence was computed as |posterior-probability-0.5|, which effectively reflects the distance of the posterior to the category boundary. To approximate the time-course of the psychophysical kernel amplitude for high and low confidence trials we varied the strength of the feedback in the model, the contrast of the orientation-selective component of the stimulus and the trial duration. The parameters used to generate the sampling model predictions were largely the same as in the original paper (κ =2, λ =3, δ =0.08, n_s_ =20, stimulus contrast on each individual frame=10, see ^17^) and only differed in the number of sensory neurons (n_x_ =256, n_g_ =64) to reduce computation time. The decreasing PKA in this model is the result of a feedback loop between the decision-making area and the sensory representation.

### Evidence-accumulation toy-model

To be able to systematically explore the predictions of attractor-based models for confidence-specific PKAs, we devised two simple abstract models. In the first the decision variable *d*_*t*_ at time *t* is defined as:

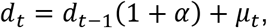

where *μ*_*t*_ is the sensory evidence at time *t*, and *α* is an acceleration parameter of accumulation process (cf. ^5^): For *α* =0 the model performs perfect integration. For *α* <0 it is a leaky integrator, and for *α* > 0 the model implements a confirmation bias or attractor. In the second model, a variant of the previous one, the acceleration parameter *α* depends on a sigmoidal function of *d* such that instead:

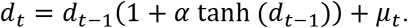

For *α* > 0 the behavior of the *d*_*t*_ can then be described by an attractor with a double-well energy landscape in which the minimum of each well correspond to the choice attractors (cf. ^16^), a behavior also observed for the sampling model by ^17^.

### Early sensory weighting model after Yates et al. (2017) ^4^

We simulated psychophysical model decisions based on sensory responses of a linear-nonlinear (LN) model. The linear stage consisted of two temporal filters (*k*, one for contrast, one for disparity):

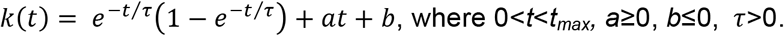

The time-varying disparity stimulus and the stimulus contrast were each convolved with the temporal filter, and their sum (*x(t)*) was exponentiated to generate spike rates:

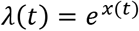

The model parameters *a*, *b*, *t*_*max*_, *τ* as well as the relative weights of the disparity and contrast kernels were chosen such that the dynamics of the output of the LN model approximately matched that of the average peri-stimulus-time histograms (PSTHs) neurons in area MT (Yates et al.; their Fig. 3b). (Starting from these initial values we then varied these model parameters to explore a range of adaptation levels as shown in supplementary Fig. 4.) To simulate the decision process we used two of these MT responses but with opposite tuning, and computed the decision variable (*d(t)*) as the integral of the difference of these time-varying MT responses. The decision on each trial was based on sign(*d(t)*) at the end of the trial, and decision confidence defined as |*d*| at the end of the trial.

To additionally account for the temporal autocorrelation of the spiking response we also simulated a variant of this basic model, also after ^4^. This variant was identical to the first except that, first, we generated spikes based on the spike rates using a Poisson process. Second, we included spike history term such that:

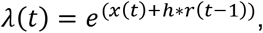

where *h* (“history filter” as in ^4^, their suppl. Fig. 1c) are the post-spike weights that integrate the neuron’s own spiking history (*r(t-1)*).

### Inclusion Criteria

Trials with fixation errors were excluded, and we only included sessions in which the animals completed at least 600 trials, and in which each experimental condition had at least 10 trials. For each session, three psychometric functions were computed (one using all the completed trials, one each including only trials for which the large available reward size was large or small, respectively). We fitted cumulative Gaussians to each of these psychometric functions, and only sessions for which each of these fits explained > 90% of the variance were included.

